# Mouse-specific but infection-unspecific IgM repertoire fingerprint following viral infection

**DOI:** 10.1101/2020.03.20.000471

**Authors:** Alexander Yermanos, Nike Julia Kräutler, Alessandro Pedrioli, Ulrike Menzel, Victor Greiff, Tanja Stadler, Annette Oxenius, Sai T. Reddy

## Abstract

Antibody repertoire sequencing provides a molecular fingerprint of current and past pathogens encountered by the immune system. Most repertoire studies in humans require measuring the B cell response in the blood, resulting in a large bias to the IgM isotype. The extent to which the circulating IgM antibody repertoire correlates to lymphoid tissue-resident B cells in the setting of viral infection remains largely uncharacterized. Therefore, we compared the IgM repertoires from both blood and bone marrow (BM) plasma cells (PCs) following acute or chronic lymphocytic choriomeningitis virus (LCMV) infection in mice. Despite previously reported serum alterations between acute and chronic infection, IgM repertoire signatures based on clonal diversity metrics, public clones, network and phylogenetic analysis were largely unable to distinguish infection cohorts. Our findings, however, revealed mouse-specific congruence between the blood and PC repertoires irrespective of infection status. Our study reveals that IgM repertoire analyses may be unsuitable for providing a fingerprint of current or previous immune challenges.

## Introduction

The possibility of personalized medicine is becoming increasingly possible due to the revolution in high-throughput sequencing (HTS) technologies (Miho *et al.*, 2018; Georgiou *et al.*, 2014; Brown *et al.*, 2019). The cost and time of sequencing an individual’s antibody repertoire has dramatically decreased over the past decade, resulting in attempts to infer disease status based on antibody repertoire sequencing (Greiff, Bhat, *et al.*, 2015). Immune-status profiling demands sufficient sensitivity and accuracy to provide correct diagnoses given the unquantifiable antigens experienced by an individual (Robinson, 2014). Immunologically intuitive metrics, such as sequence diversity, clonal expansion, and germline gene usage have been routinely employed to quantify antibody repertoire fingerprints between different vaccine and infection conditions, based entirely on the antibody repertoire (Jackson *et al.*, 2014; Jiang *et al.*, 2013; Greiff *et al.*, 2017). In human patients, however, most antibody repertoire sequencing experiments are limited to circulating B cells in the peripheral blood (Jackson *et al.*, 2014; Doria-Rose *et al.*, 2014; Wu *et al.*, 2015; Tsioris *et al.*, 2015; Vander Heiden *et al.*, 2017). This implicitly enables time-resolved sampling of the antibody repertoire within the same host over time, despite sacrificing spatial and physiological resolution from repertoires across multiple organs. Furthermore, peripheral blood heavily biases the cellular composition to naïve B cells of the IgM isotype, as seen with single-cell sequencing experiments (Horns *et al.*, 2020). While previous studies have described and classified infection status based on antibody repertoire sequencing (Greiff, Bhat, *et al.*, 2015; Emerson *et al.*, 2017), it remains largely unknown how multiple sampling time points, antibody isotype, and organ selection impacts these fingerprints, especially in the context of viral infection.

To quantify whether the aforementioned parameters can distinguish viral infection cohorts, we utilized both temporally- and spatially-resolved antibody repertoire sequencing data from mice infected with lymphocytic choriomeningitis virus (LCMV) (Kräutler *et al.*, 2020). LCMV is a rodent-borne pathogen that can elicit either an acute (resolved within weeks) or chronic (resolved within months) infection depending on the initial viral strain and dose. It has been demonstrated that CD8 T cells are necessary for the clearance of acute LCMV infection, whereas the conversion to a follicular response is crucial to resolve persisting LCMV infection via virus-neutralizing antibodies (Planz *et al.*, 1997; Thomsen *et al.*, 1996; Greczmiel *et al.*, 2017). Although both B and CD4+ T cells are dispensable for the resolution of acute LCMV infection, an increase in both IgG and IgM titers against the purified virus has nevertheless been observed in both infection cohorts (Kräutler *et al.*, 2020). Despite this increase in serum titers against purified virus for both isotypes, the IgG isotype (particularly IgG2c) has been shown to be crucial to resolving persistent LCMV infection (Barnett *et al.*, 2016). It has, however, also been demonstrated that the early IgM response can influence the clearance of chronic LCMV infection in the context of transgenic mice expressing virus-neutralizing antibodies (Seiler *et al.*, 1998). While viral specific fingerprints in the IgG repertoire have been observed following acute and chronic LCMV infection (Kräutler *et al.*, 2020), it remains unknown whether this holds similarly true for the IgM repertoire.

Therefore, we employed a bioinformatic framework to quantitatively characterize the IgM antibody repertoire following acute and chronic LCMV infections. Our analysis leveraged metrics quantifying clonal expansion, germline gene usage and the extent of clonal convergence across and within IgM repertoires. We discovered that both acute and chronic LCMV infection had minimal effects on the clonal composition of the IgM B cell repertoire compared to uninfected mice. While cohort-specific IgM repertoire signatures were minor, mouse-specific repertoires showed high congruence between the blood and PC compartments, irrespective of infection cohort. Compared to IgG repertoires, our findings reveal a severe limitation of IgM repertoire analyses in providing a fingerprint of actual or previous immune challenges.

## Results

### Minor influence of LCMV infection on IgM clonal expansion

We utilized bulk antibody heavy chain repertoire sequencing from a previously published experiment in which repertoires were sequenced longitudinally 10 days before infection and 10, 20, 50, 60 and 70 days post infection (dpi) for 15 animals (n_naive_=5, n_acute_=5, n_chronic_=5), with the exception of two time points (one sample 10 days before infection and one sample 60 dpi) due to failed library amplification (Kräutler *et al.*, 2020). This dataset furthermore includes bone marrow plasma cells (BM PCs) that were sorted by flow cytometry (FACS) 70 dpi for 14 of the 15 mice and amplified using IgM and IgG specific primers. We focused our analysis on the top 1,000 IgM clones for each repertoire to avoid variations in sequencing depth and to ensure that all samples were compared using the same number clones.

We first asked whether viral infection induced clonal expansion of the circulating blood IgM repertoire shortly after infection. We initially visualized the clonal frequency (percent of top 1,000 clones) for the top 50 clones either before infection (−10 dpi) or 10 dpi for an exemplary mouse from each cohort (Figure 1A). We immediately observed that the clonal abundance of the most expanded IgM clone increased from approximately 0.5% to 1.5% of the top 1,000 clones within individually chronically infected mice but not in the other two cohorts. Quantifying the ratio of the most abundant clone from either −10 dpi to 10 dpi in each animal revealed the trend that the frequency of the top clone in infected animals doubled, with the effect more apparent in the chronically infected cohort (Figure 1B). To further quantify the clonal expansion for all IgM clones, we calculated the commonly utilized Shannon evenness (Greiff, Miho, *et al.*, 2015) for each blood repertoire at each time point (Figure 1C). A Shannon evenness of 1 indicates a homogenous degree of clonal expansion across all clones within the repertoire, whereas a value of 0 signifies varying degrees of clonal expansion. Consistent with the expansion of the top IgM clone in chronically infected mice, the Shannon evenness in this cohort was lower than the other two cohorts 10 and 20 dpi (Figure 1C). Interestingly, the Shannon evenness was low 70 dpi for all three cohorts, potentially due to the increased volume of sampled blood at the terminal time point (Kräutler *et al.*, 2020). The Shannon evenness of the IgM BM PC repertoire was comparable for all three cohorts (Figure 1D), with much lower values when compared to the blood repertoires (Figure 1C), indicating that IgM-producing PCs in the BM were enriched in expanded clones, presumably due to focusing on antigen-experienced cells. We next questioned if the highly expanded clones at early time points remained expanded at subsequent time points. To quantify this, we calculated the Spearman correlation of the clonal frequencies of clones found at both 10 dpi in blood and in the BM PC repertoire 70 dpi for a single chronically infected mouse (Figures 1E). Quantifying this for each mouse across all cohorts revealed minimal correlation between clonal expansion of clones found at 10 dpi and in the BM PC 70 dpi (Figure 1F). Performing the same analysis for clones observed at both −10 and 10 dpi revealed higher clonal frequency correlation coefficients for each cohort compared to the previous comparison, suggesting that the time between sampling, and not viral infection, is the major driver of correlated clonal expansion (Figure 1G). Indeed, comparable correlation coefficients were also observed when comparing blood repertoires before or shortly after viral infection (Figure 1H), further highlighting the minor impact of viral infection on the IgM repertoire. Taken together, these data suggest that chronic but not acute infection impacted, but only to a minor extent, the clonal expansion profile of highly expanded clones at an early time point after infection.

**Figure 1.**
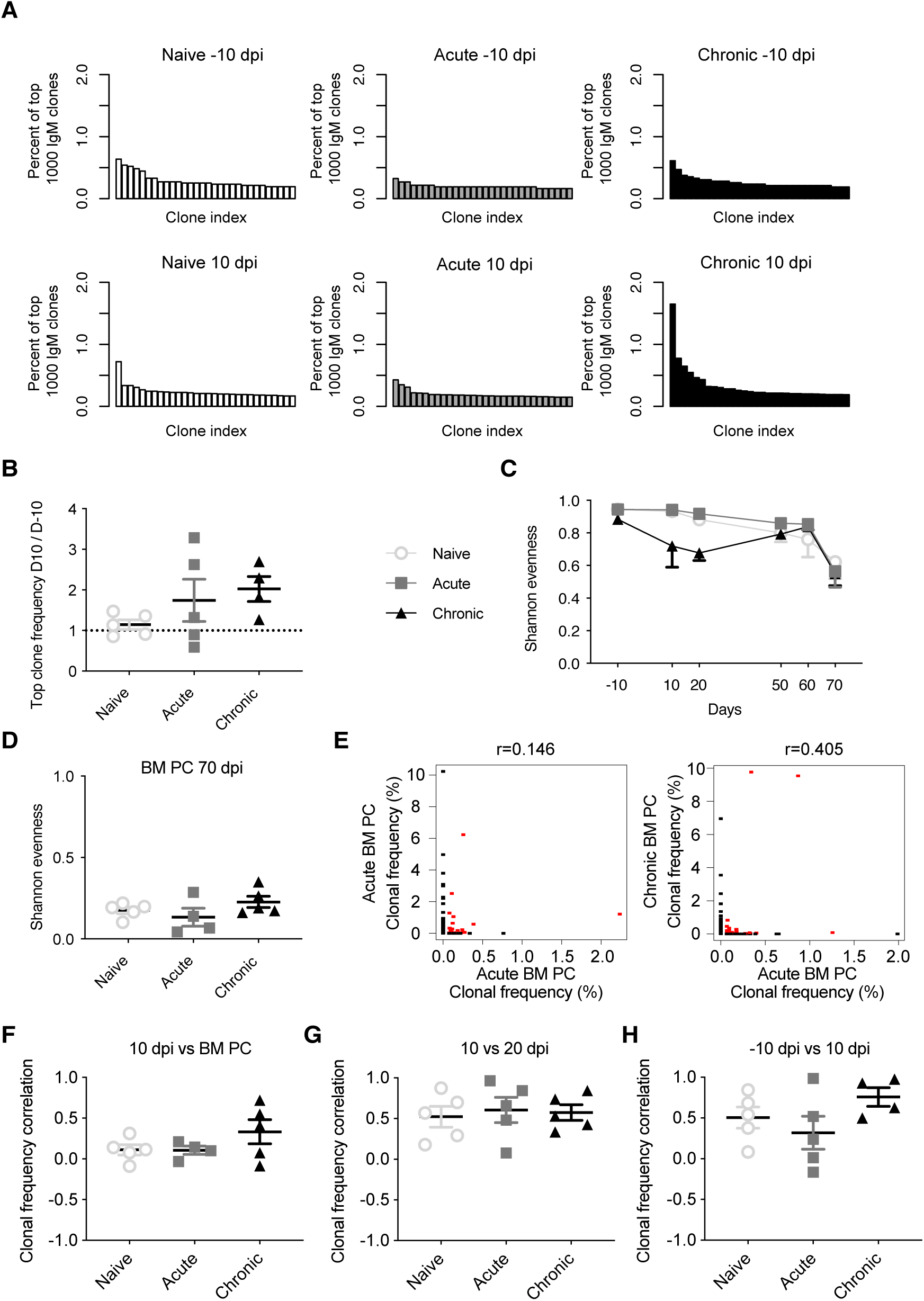
Time-resolved clonality of the IgM repertoire following acute and chronic LCMV infection. (A) Exemplary clonal frequency distributions of the most abundant IgM clones for the indicated time points and infection cohorts. (B) Ratio of the clonal frequency (% of IgM clones) of the most expanded IgM clone within the same mouse either 10 days post infection (dpi) or 10 days before infection (−10 dpi). (C) Shannon evenness quantifying clonal expansion of blood repertoires over time. D) Shannon evenness quantifying clonal expansion of bone marrow (BM) plasma cells (PCs) 70 dpi. (E) Example Spearman correlation of clonal frequencies for those clones found at either 10 dpi and in BM PC repertoire of the same mouse. Red points indicate identical clones found at both time points, whereas black points indicate clones found in only one time point. (F) Spearman correlation coefficients describing clonal frequency of those clones found in both blood 10 dpi and BM PC repertoires of the same mouse. (G) Spearman correlation coefficients describing clonal frequency of those clones found in blood repertoires both 10 and 20 dpi in the same mouse. (G) Spearman correlation coefficients describing clonal frequency of those clones found in blood repertoires both −10 and 10 dpi in the same mouse.

### Distinct V gene usage between blood and BM PC repertoires

After observing minor differences between the clonal frequencies of uninfected and infected animals (Figure 1), we asked whether LCMV infection resulted in the recruitment of B cells with specific patterns of germline gene usage. To answer this question, we calculated the percent of unique clones using each V gene for both blood and BM PC repertoires. As an example, we plotted the most utilized V genes for blood 10 dpi and the BM PC repertoires for a single chronically infected mouse (Figure 2A). We observed a majority of clones using V14-2, V14-4, and V5-17 in the blood repertoires (>30% of clones), whereas these V genes accounted for a smaller fraction in the BM PC compartment of the same animal (<10%) (Figure 2A). Quantifying this for each cohort across all time points revealed consistent V gene patterns differentiating blood and BM PC repertoires, with the aforementioned V genes remaining highly expressed in the blood repertoires throughout the experiment in all cohorts (Figure 2B). Furthermore, the BM PC repertoires for all infection groups showed more diverse V gene usage, with the median usage for each V gene higher than in the blood repertoires (Figures 2B, 2C). While the most striking difference in V gene usage was between the repertoires of the different organs, there were nevertheless trends of cohort specific fingerprints. We observed the trend that certain V genes were upregulated following acute LCMV infection, as exemplified by an increased proportion of clones using V1-72 10 dpi in acutely infected animals but not in the other cohorts (Figure 2D). We finally asked if viral infection resulted in differential expression of V genes when comparing blood repertoires −10 dpi to 10 dpi. Quantifying the log_2_-fold change for all V genes uncovered signatures of differential germline expression between these two time points (Figure 2E). However, this effect was minimal for both cohorts relative to those differences between the blood and BM, with a much wider range of up- or down-regulation for V genes in the BM PC repertoires (Figure 2E). These findings suggest that viral infection has relatively little influence on the clonal composition of the most abundant IgM clones. Furthermore, blood and BM PC repertoires showed distinct patterns of V gene usage, with the BM PC repertoires consisting of a more diverse set of V genes for all cohorts.

**Figure 2.**
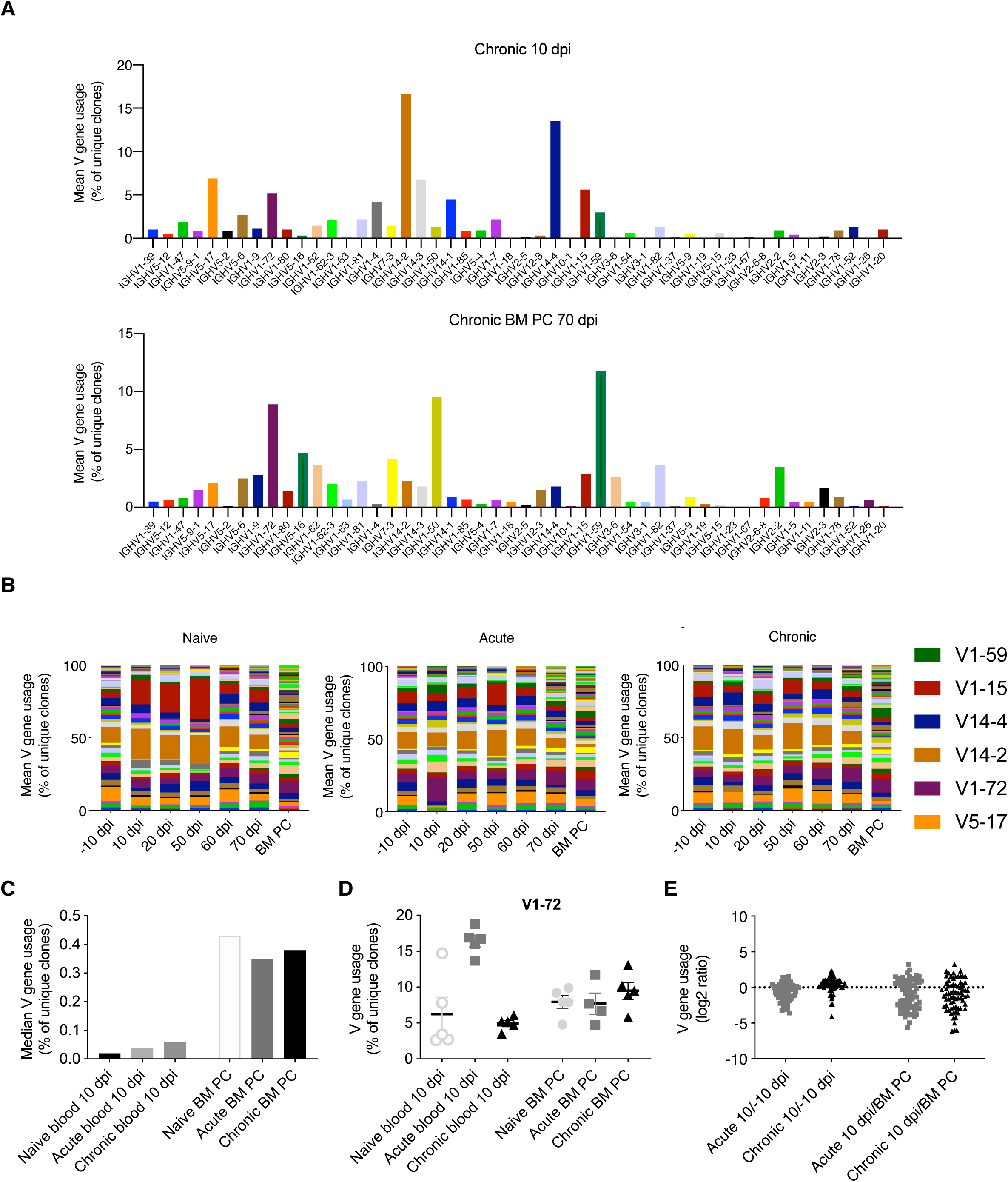
Distinct germline gene usage in blood and BM PC repertoires. (A) Percent of unique IgM clones using the indicated subset of IgH V gene in a single chronically infected mouse at either the blood repertoire 10 dpi (top) or in the bone marrow plasma cell repertoire 70 dpi (bottom). (B) The average percent of unique IgM clones using a given V gene for each cohort. Colors correspond to distinct V genes and the value corresponds to the percentage of usage. Selected genes are highlighted. (C) Median V gene usage across all V genes for each cohort. (D) The percent of unique IgM clones using IgH V1-72 in the blood repertoire 10 dpi or in the BM PC compartment 70 dpi. (E) The log_2_ ratio quantifying the up- or down-regulation of a given V gene at the indicated time points. Each point corresponds to a single V gene. Positive values indicate increased usage across the cohort in the blood repertoire 10 dpi compared to either −10 dpi (left) or BM PC 70 dpi (right).

### Public clones do not account for compartment-specific V gene usage

After observing consistent V gene usage in the blood and BM PC repertoires across all cohorts, we next asked if this was due to a large proportion of public clones, defined as clones with identical CDRH3 sequences (amino acid) found in at least two mice. Quantifying the percent of clones found in multiple mice revealed that both the pooled blood and the BM PC repertoires were largely private, with less than 7% of clones found within multiple mice (Figure 3A). Despite the low incidence of public clones, we were curious as to whether these clones employed a distinct repertoire of V genes. To that end, we quantified the percentage of clones using a given V gene for both the public and private (clones found in only one mouse) clones in the pooled blood repertoires and the BM PC repertoires (Figure 3B). Despite their low frequency, the public clones showed a biased V gene usage relative to the private clones in both blood and BM PC repertoires, with certain V genes, such as V14-2 and V12-3, more present than in the corresponding private repertoires (Figure 3B). Similarly, there was a trend for shorter CDRH3 lengths in the public clones compared to the private clones for both compartments, further suggesting possible selective pressures for public clones (Figure 3C). Given these aforementioned properties specific for public clones, we wondered if the majority of the public clones occurred in the groups exposed to viral infection. Therefore, we calculated the pairwise Jaccard index, a metric ranging between 0 and 1, quantifying the degree of overlap between two samples. Similar to the previously described percentage of unique clones (Figure 3A), the pairwise overlap for both blood and plasma cell repertoires was low, with Jaccard indices less than 0.05 for all animals, corresponding to ∼5% overlapping clones (Figures 3D, 3E). While there was a trend of acutely infected animals displaying lower Jaccard overlaps in the pooled blood repertoires, all cohorts nevertheless showed higher convergence than the accompanying BM PC repertoires (Figures 3D, 3E). Together, these data suggest public IgM clones arise in a viral-independent manner and the majority of blood and BM PC clones are primarily confined to a single mouse.

**Figure 3.**
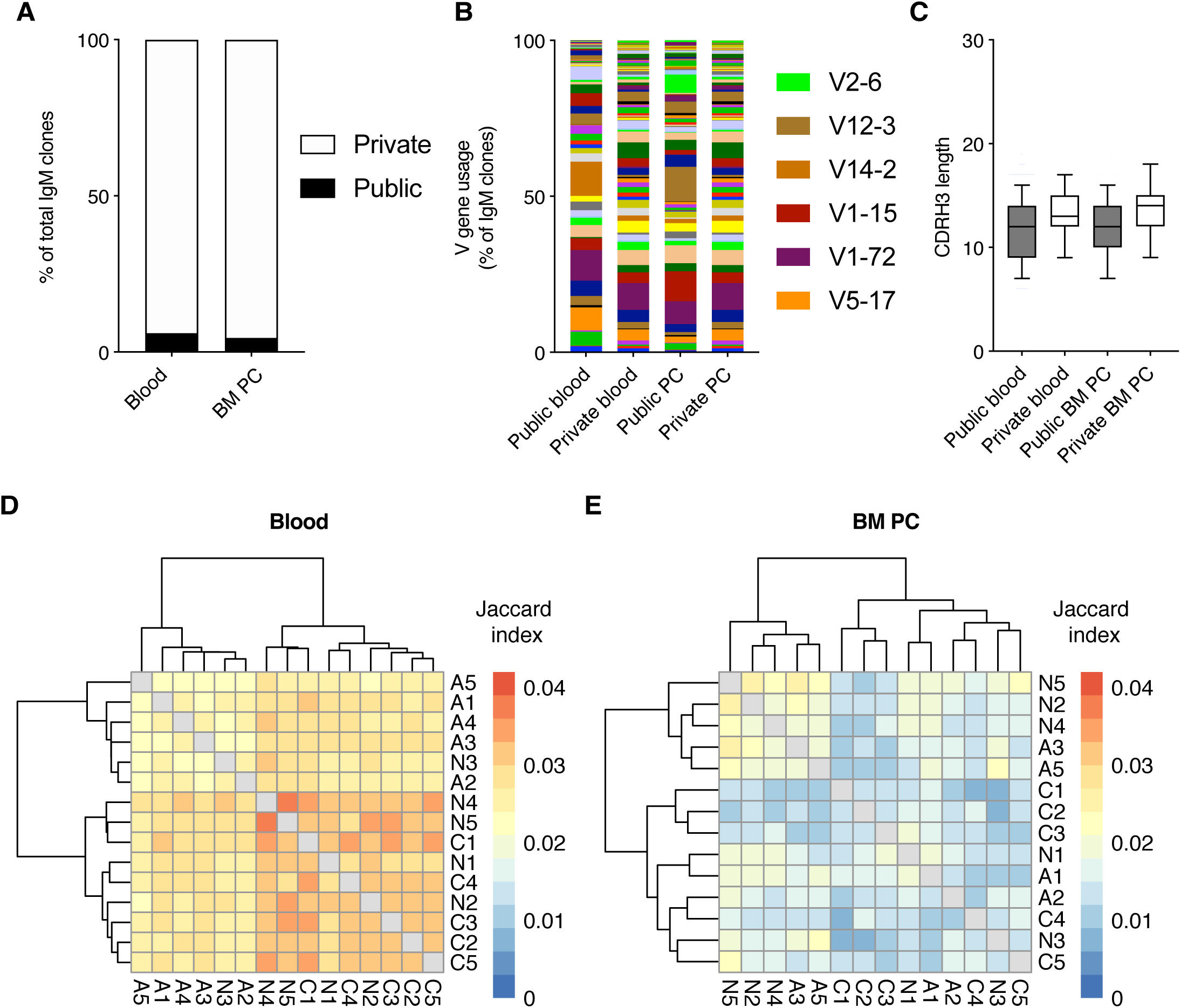
Minor but distinct profile of public clones across all infection cohorts. (A) The percentage of pooled blood or BM PC IgM clones either private (found in only 1 mouse) or public (found in more than one mouse). (B) V gene usage of public clones and private IgM clones from pooled blood and BM PC repertoires. (C) Amino acid CDRH3 length of public and private clones from pooled blood and BM PC repertoires. (D) Pairwise Jaccard index quantifying clones found in the pooled blood repertoires of the indicated two mice. Labels correspond to infection cohort (N=uninfected, A=acute infection, C=chronic infection) and number corresponds to a distinct mouse within the cohort. Intensity corresponds to the number of clones common to both repertoires divided by the number of unique clones found in either repertoires. Sample order and accompanying dendrograms were determined via hierarchical clustering. (E) Pairwise Jaccard index as in D but quantifying clones found in the BM PC repertoires of multiple mice.

### Mouse-specific overlap of blood and BM PC repertoires

Both the large discordance of V gene usage for all cohorts (Figure 2) and the low frequency of public clones (Figure 3) suggested that only a small fraction of clones would be shared between blood and BM PC repertoires. Contrary to this expectation, a large percentage (>10% for all cohorts) of the clones from the BM PC repertoire had been previously sampled in an earlier blood time point (Figures 4A, 4B). This effect, however, was not cohort-specific, indicating that there is a general overlap between the circulating IgM B cells and the BM PC repertoires. This increased overlap was mouse-specific, but not cohort-specific, indicated by the lower clonal overlap of the blood and BM PC repertoires from two distinct animals within the same infection group (Figures 4A, 4B). We next asked whether the sampling time impacted the extent of the clonal overlap between the two compartments. On average, ∼25 clones of the PC repertoire were also present in the same animal 10 days before infection, regardless of cohort (Figure 4C). Again, this trend was mouse-specific, as the overlap between the blood repertoire −10 dpi with the BM PC across different animals was lower than within the same animal (Figure 3C). In comparison, the blood and BM PC repertoires shared a larger number of clones when sampled at the same time point for all three cohorts (Figure 4D). We next asked whether the clones shared between the blood and BM PC repertoires were composed of similar germline elements for all cohorts. V gene usage profiles between the PC clones also in the blood to those clones private to the BM PC repertoire were similar across all animals, with certain V genes such as V12-2, V1-72, and V1-80 being utilized in all mice (Figure 4E). While some V genes were expressed at higher levels in individual animals (e.g. V1-50 and V5-4), this effect was largely cohort-unspecific, further suggesting a minor impact of viral infection on clonal convergence between the blood and BM PC IgM repertoires. We lastly asked whether LCMV resulted in an increase of persisting clones (identical clones found within a given mouse at multiple time points) in the blood repertoires early after infection. There were minor differences between the three cohorts, with the clonal persistence for all cohorts ranging from ∼1 to 10%, further suggesting a limited role of LCMV infection on both the blood and BM PC IgM repertoire.

**Figure 4.**
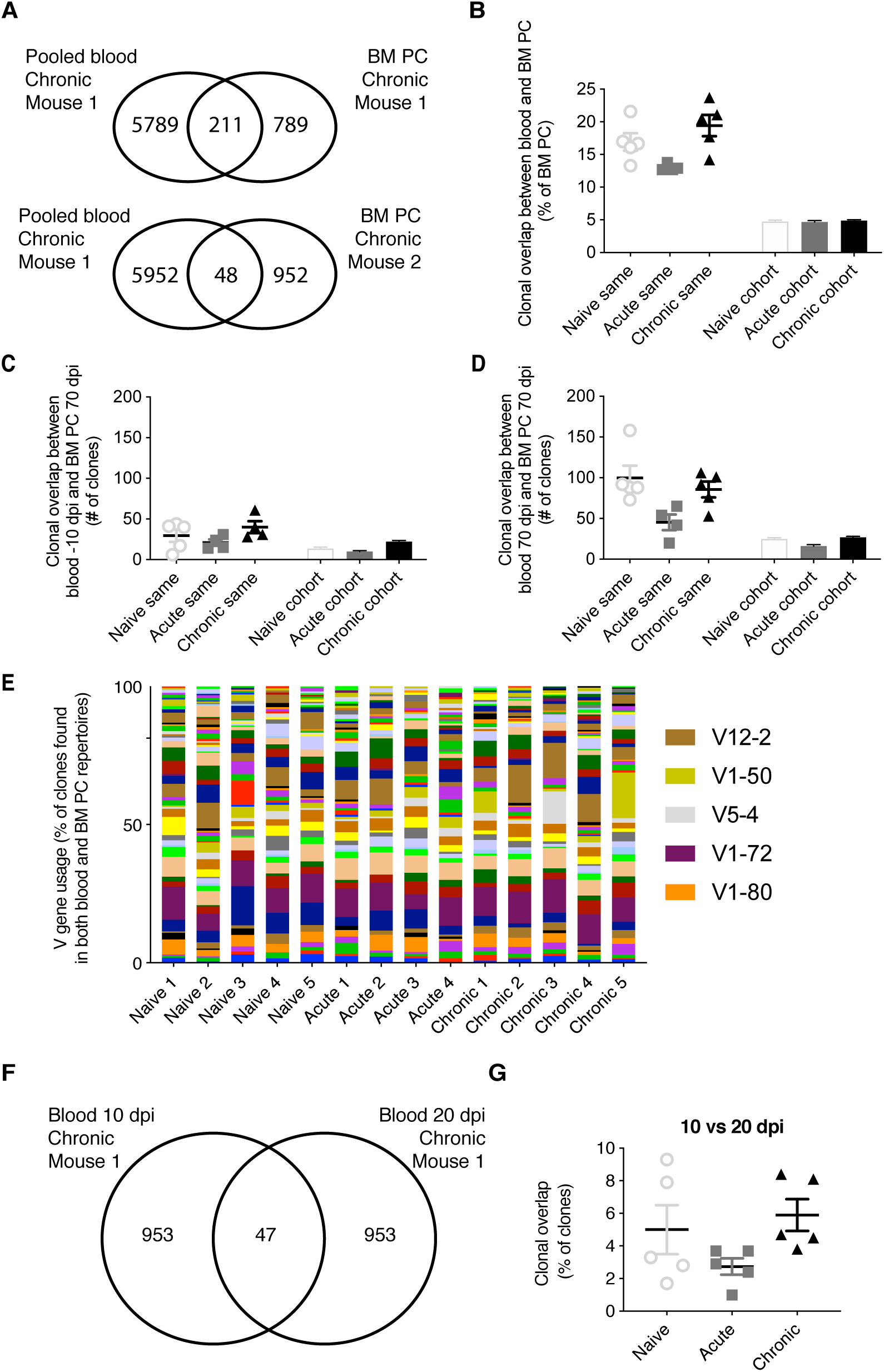
Mouse-specific overlap between blood and bone marrow plasma cell repertoires. (A) Example clonal overlap between the pooled blood repertoires with the BM PC repertoire of the same mouse (top) or a different mouse (bottom). (B) The percent of IgM BM PC clones present in the blood repertoire of either the same mouse or other mice of the same cohort. (C) The number IgM clones present in both the blood 10 dpi and BM PC 70 dpi in either the same mouse or other mice of the same cohort. (D) The number IgM clones present in both the blood 70 dpi and BM PC 70 dpi in either the same mouse or other mice within the same cohort. (E) V gene usage of those clones found in both the blood and BM PC repertoires. (F) Example overlap of clones found in both the blood IgM repertoires 10 and 20 dpi of a single chronically infected mouse. (G) Clonal overlap between the blood IgM repertoires for the same mouse at 10 or 20 dpi.

### Minor signatures of clonal convergence induced by LCMV infection

Until now, our analysis had relied on describing the antibody repertoire based on identical CDRH3 amino acid sequences, which inherently ignores any relationship between highly similar antibody sequences. This led us to next ask whether patterns of convergent selection following LCMV infection could be detected if we examined the presence of closely-related antibody sequences. To this end, we first quantified the mean pairwise edit distance, which quantifies the number of amino acid mutations separating two sequences, for each mouse at each time point. The edit distance remained constant throughout the 80 days for all cohorts in both blood and BM PC repertoires (Figures 5A, 5B), suggesting that IgM repertoires remained globally unchanged for all animals. We next questioned if focusing the analysis on those highly similar clones would elucidate differences between the infection cohorts. Thus, we performed an in-depth sequence similarity analysis to construct networks (Miho *et al.*, 2018) for the IgM repertoires within each mouse, with edges indicating unique clones separated by less than 3 amino acid mutations (Figure 5C). As an example, we plotted the similarity networks for the blood repertoires 10 dpi for two infected mice, revealing both connected and unconnected clones for each mouse (Figure 5C). To formally quantify this across all time points and mice, we quantified the clustering coefficient for each network, a metric describing the extent of connections between vertices. This analysis again revealed no differences between infected and uninfected animals in both the blood and BM PC repertoires (Figures 5D, 5E). We next questioned asked if convergent selection would be observed when incorporating the time-resolved component into the analysis. We thus pooled all clones from the blood and BM PC repertoires for each mouse and again calculated similarity networks for each cohort. While the clustering coefficient remained comparable for all cohorts after pooling all time points (Figure 5F), we indeed observed a reduced network density (ratio of connections to total possible connections) of acutely infected mice relative to the other two cohorts (Figure 5G), but comparable network diameters (the longest path of connected clones). While minor differences may represent subtle changes to the overall sequence space of the IgM repertoire, the majority of these analyses further imply that LCMV infection has a minor influence on the clonal selection of IgM blood and BM PC repertoires.

**Figure 5.**
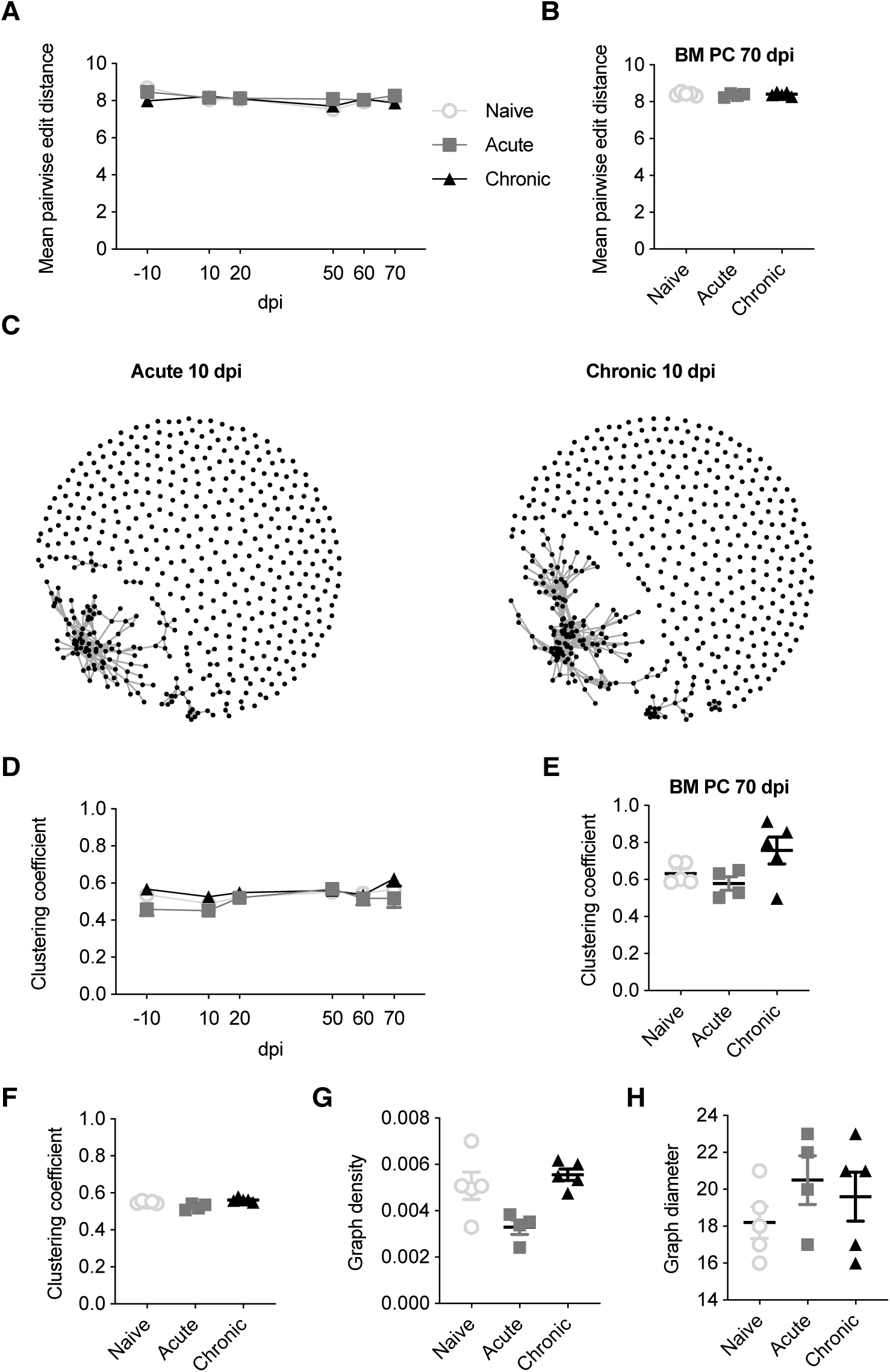
Sequence similarity metrics do not differentiate infection cohorts. (A) The mean pairwise edit distance between all clones for each mouse either in the blood or (B) BM PC IgM repertoires. (C) Example similarity networks of naïve, acute and chronic IgM clones 10 dpi. Vertices indicate a unique CDRH3 clone. Edges indicate CDRH3 sequences separated by less than three amino acid mutations. (D) Clustering coefficient of the similarity networks for the blood repertoire at the indicated time point for each mouse. (E) Clustering coefficient of the similarity network for the BM PC repertoire of each mouse. (F) Clustering coefficient of the similarity networks for the pooled IgM clones across all time points and compartments for each mouse. (G) Graph density of the similarity networks for the pooled IgM clones across all time points and compartments for each mouse. (H) Graph diameter of the similarity networks for the pooled IgM clones across all time points and compartments for each mouse.

### Mouse- and cohort-specific overlap of class-switched clones

We finally questioned asked how frequent class-switched IgG clones were detected with identical IgM clones within the same repertoire. We thereby calculated the percentage of IgM clones with an identical IgG CDRH3 sequence in the blood repertoire for each mouse across the 80 days (Figure 6A). This quantification revealed an increase in the overlap of IgG and IgM clones following both acute and chronic infection 10 dpi. This increase was maintained throughout the remainder of the chronic infection, whereas the overlap of acutely infected animals returned to levels observed in uninfected animals (Figure 6A). While immediately striking, this increased overlap between IgG and IgM sequences in the infected cohorts was likely due to the increased number of unique clones in the IgG repertoires induced by chronic and acute infections (Kräutler *et al.*, 2020), as the cohort-specific effect was reduced after normalizing by the percent of IgG observed in the IgM compartment (Figure 6B). A similar phenomenon was observed in the BM PC repertoires, with increased convergence of IgG and IgM clones when normalizing by the amount of IgM but not when normalizing by the amount of IgG (Figures 6C, 6D). We wondered, however, if a stronger convergence would be observed when pooling all repertoires (both blood and BM PC) for each mouse and then again quantifying the convergence between IgG and IgM clones. Indeed, we observed a trend that the overlap between IgG and IgM was higher in both acute and chronic cohorts, even after normalizing by either isotype (Figure 6E). Furthermore, we observed that the overlap between the pooled repertoires of a given mouse was higher than two distinct mice within the same cohort (Figure 6E). Accompanying this realization, we noticed a higher degree of clonal convergence within both the acute and chronic cohorts compared to the naive cohort, indicated by the higher Jaccard indices when comparing the overlap between IgG clones of one infected mouse to the IgM clones another (Figure 6E). Together, these findings reveal minor patterns of convergence between the IgG and IgM repertoires of infected mice, and that this convergence is again mouse-specific.

**Figure 6.**
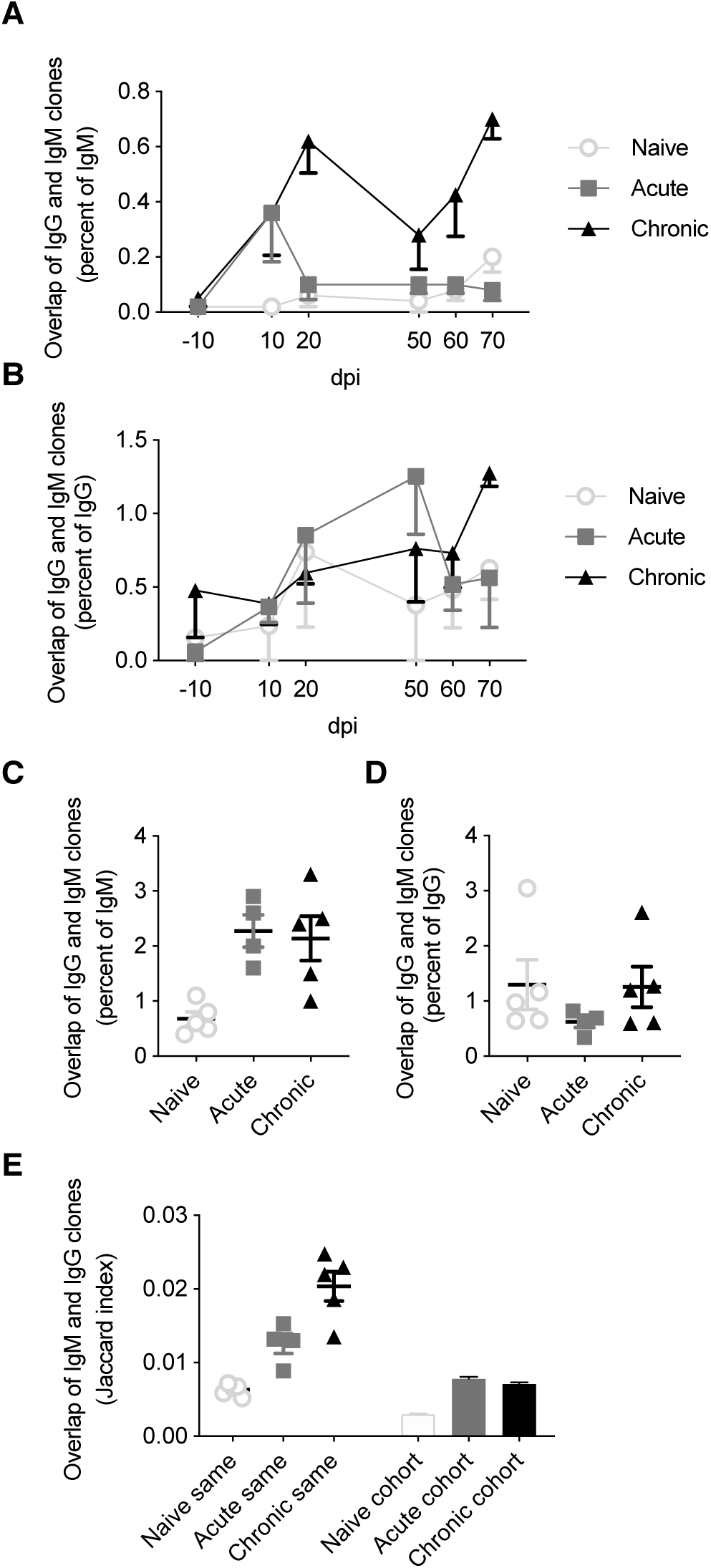
Increased overlap between IgG and IgM repertoires following LCMV infection. (A) The percentage of IgM clones with a corresponding IgG clone in the same blood repertoire at the indicated time point. (B) The percentage of IgG clones with a corresponding IgM clone in the same blood repertoire at the indicated time point. (C) The percentage of identical IgM clones observed in the IgG repertoire of the same BM PC sample. (D) The percentage of identical IgG clones observed in the IgM repertoire of the same BM PC sample. (E) Jaccard index quantifying overlap between the pooled IgM and IgG repertoires of either the same mouse or of different mice within each cohort.

## Discussion

As the amount of high-throughput antibody repertoire sequencing data increases, attention is being drawn towards the possibility of using information extracted from repertoires as a possible immunodiagnostic. In the context of human patients, this would rely upon sampling circulating lymphocytes in the blood, subsequently sequencing antibody repertoires, and finally deconvoluting immune histories in a potentially pathogen specific manner. As it is difficult to connect clonal information from circulating repertoires to tissue-resident B cells, significant weight is given to the snapshot presented by the blood repertoire. Here we have leveraged an extensively-studied murine viral infection model that allowed for both the deconstruction of antibody repertoires in both circulating and tissue-resident B cells. It has been previously described that cohort-specific signatures arise in the IgG repertoires following LCMV infection, in addition to a higher level of congruence between the blood and BM PC IgG repertoires following chronic LCMV infection relative to acutely-infected and uninfected animals (Kräutler *et al.*, 2020). Given the progression of class switching from IgM to IgG isotype following the interaction between the BCR and its cognate antigen, we hypothesized that the cohort specific signatures previously observed in the IgG repertoire would be similarly present in the most expanded portion of the IgM repertoires.

Surprisingly, multiple metrics revealed minimal differences in IgM repertoires of uninfected and infected mice, both in blood repertoires sampled across six different time points and in the BM PC compartment. One of the few metrics distinguishing the IgM repertoires of the three included cohorts was the clonal expansion profile, where increased clonal expansion at 10 and 20 dpi was observed in chronically infected mice (Figure 1). Both the importance and stability of these expanded clones remains questionable, however, as very few clones (<10%) retained in the blood repertoires from 10 to 20 dpi for all cohorts, irrespective of infection history (Figure 4F). Additionally, the degree of clonal expansion of the clones present at both 10 dpi and 20 dpi was mildly correlated for all infection cohorts, further suggesting that this degree of clonal persistence is an inherent property of the blood repertoire over time. Other metrics, such as V gene usage, public clones, and network analysis revealed little to no differences between the infection cohorts (Figures 2, 3, 5). In a few analyses, such as the network density (Figure 5G) and public clones in the blood repertoires (Figure 3C), acutely infected animals showed distinct patterns of selection after pooling clones from all time points. While these differences were nevertheless subtle, they may arise due to the recruitment of some LCMV specific B cells shortly after infection that do not remain circulating in the blood after viral clearance by 20 dpi. This hypothesis is partially supported by previous work which observed an increase in unique IgG clones in the blood 10 dpi following chronic and acute LCMV infection that only maintained at later time points in the chronically infected cohort (Kräutler *et al.*, 2020). This does not necessarily explain, however, why certain V genes (such as V1-72) were much more present 10 dpi in the acute but not the chronically infected cohort (Figure 2D).

The investigated dataset further allowed for a comprehensive comparison of blood and BM PC repertoires of the same mice, in both infected and uninfected repertoires. One consistent finding was that distinct patterns of V gene usage were observed when comparing the blood and BM PC repertoires, consistent with previous reports describing distinct organ-specific repertoires (Figure 2) (Briney *et al.*, 2014; Greiff *et al.*, 2017). The less diverse germline usage of the BM PC repertoires could reflect preferential selection and specificity of certain V genes recruited to the bone marrow niche, whereas the patterns observed in the blood repertoire may reflect the underlying probability distribution of a certain V gene to be selected during VDJ recombination, as previous works have described (Elhanati *et al.*, 2014; Marcou *et al.*, 2018). While the characterization of public clones and sequence similarity networks revealed similar degrees of clonal convergence of the IgM response across all three cohorts, we discovered a strong mouse-specific convergence between the blood and BM PC IgM repertoires, with ∼15% of BM PC clones found in the earlier IgM time points (Figure 4B). Surprisingly, this convergence was apparent when comparing the clonal overlap between blood and BM PC repertoires of the same mouse but not the cohort up to 80 days before sacrifice (Figure 4C), further highlighting the mouse-specific nature of the IgM repertoire.

It is possible that focusing the analysis upon the virus-specific IgM repertoire would reveal greater differences between acute and chronic viral infection, however, the comparable V gene usage, quantity of public and persisting clones, and clonal convergence between the three cohorts suggests otherwise. Another limitation from the analyzed data set is the lack of light chain information, as it has previously been observed that clonal expansion and V gene usage can be altered when light chain information is included in the analysis (DeKosky *et al.*, 2015). While technological advancements in single-cell sequencing currently provide both pairing information and reliable clonal frequency profiles that can be easily validated for specificity (Horns *et al.*, 2020), it is questionable whether this increased information would be useful for *in silico* classification of disease status based on IgM repertoire sequencing (Zhou and Kleinstein, 2019). Furthermore, current single-cell sequencing techniques are often limited to much lower depth (thousands of clones) compared to bulk sequencing (millions of clones), which suggests that similar results would be obtained from these pipelines. In summary, our findings demonstrate inherent differences between the blood and BM PC IgM repertoires and that these immune profiles are rather robust even to high-dose LCMV infection. Our findings, in conjunction with previously described infection-specific IgG fingerprints (Kräutler *et al.*, 2020), casts doubt upon the usefulness of non-antigen-specific heavy chain IgM repertoire sequencing to distinguish past viral immune responses.

## Methods

### Repertoire analysis

Raw sequencing data was taken from accession number E-MTAB-8585 (Kräutler *et al.*, 2020) and aligned to the murine germline segments using MiXCR (v2.1.1) with clones defined as unique CDRH3 amino acid for all analyses (Bolotin *et al.*, 2015), as previously reported (Greiff *et al.*, 2017). Sequences aligning to either IgM or IgG were retained for further downstream processing. The top 1,000 IgM clones based on clone count were included for each of the sequencing files. Germline gene usage for each clone was determined by the germline segment with the top alignment score determined by MiXCR. Clonal frequency was calculated as the MiXCR-determined clone count divided by the total clone count of the included IgM clones. Top clonal frequency was calculated as the percentage of the top 1,000 clones for the clone with the highest clone count found 10 dpi divided by the percentage of the top 1,000 clones for the clone with the highest clone count found −10 dpi for each mouse. The Shannon evenness was calculated by taking the exponential of the diversity function in R package vegan (Oksanen *et al.*, 2019) subsequently dividing by the number of clones for each repertoire, as previously described (Greiff, Bhat, *et al.*, 2015). V gene usage was always calculated as the percent of unique clones, thereby ignoring the clone count component as previously reported (Greiff *et al.*, 2017). Log_2_ ratios resulted from dividing the mean V gene usage of all mice within a cohort by the mean V gene usage across all animals in the second cohort.

Public clones were defined as any amino acid CDRH3 sequence found in more than one mouse in either the blood or BM PC, respectively (Kräutler *et al.*, 2020). Jaccard indices were calculated by quantifying the intersection of CDRH3s between two samples divided by the length of the union of the same samples. Clonal overlap was always calculated based on identical amino acid sequences between any two repertoires. Edit distance was calculated using the function stringdistmatrix from the R package stringdist (v0.9.5.5) (Loo, 2014) with the method set to “lv”.

Similarity networks were produced by first inferring an adjacency matrix based on the output from stringdistmatrix and subsequently setting all values separated by less than 3 amino acid sequences to 1. All other values were set to 0, and the remaining matrix was used as input for the function graph_from_adjacency_matrix in the R package igraph (v1.2.4.1) (Csardi and Nepusz, 2006) with mode set to undirected. Clustering coefficients, graph density, and diameter were calculated for all networks using the functions transititivity, graph.density, and diameter in igraph, respectively, with the type parameter set to “global”. Heatmaps were created using the Rpackage pheatmap, with arguments cluster_rows and cluster_cols set to true. All other plots were produced in Graphpad prism v8. All error bars indicate the standard error of mean.

